# Metabarcoding reveals that bacterial and fungal microbiomes are perturbed by micropollutants in a fjord system (Hakefjorden and Askeröfjorden) at the Swedish west coast

**DOI:** 10.64898/2026.03.13.711609

**Authors:** Eduardo Acosta, Nicolai Verbücheln, Sonja Schaufelberger, R. Henrik Nilsson, Werner Brack, Alexis Fonseca, Thomas Backhaus, Pedro A. Inostroza

## Abstract

Fjord systems are susceptible to anthropogenic pressures, including discharges from wastewater treatment plants (WWTPs), which introduce micropollutants into coastal waters. We investigated the impact of micropollutants on bacteria and fungi within a fjord system adjacent to a significant petrochemical industry hub on the Swedish west coast. We characterised microbial assemblages along a land-to-sea transect, encompassing freshwater streams receiving agricultural and urban runoff, as well as the direct effluent from a WWTP. Our findings revealed elevated concentrations and a diverse array of micropollutants in the WWTP effluent and the stream running through the urban/industrial zone, highlighting these areas as major sources of pollution to the fjord. Bacterial and fungal communities inhabiting the WWTP effluent and the receiving marine waters near the marine outflow exhibited distinct structural compositions, indicating a selective pressure exerted in part by the micropollutant load. While freshwater sites generally displayed higher overall microbial diversity compared to marine sites, the WWTP effluent showed reduced diversity in both bacterial and fungal communities, likely due to the impact of micropollutants. Interestingly, marine sites far from the WWTP discharges exhibited a recovery in bacterial diversity, suggesting a potential response or adaptation. In contrast, fungal diversity remained comparable to that observed in other marine locations. Multivariate analyses identified physicochemical parameters and nutrients, alongside with summed fungicides and antibiotic stress as key factors driving the community dissimilarities across the fjord. Significant disruptions in potential bacterial metabolism and fungal ecological functions were evident at the WWTP discharge point, underscoring the ecological consequences of wastewater pollution.

**Highlights:**

- WWTP discharge is the primary source of complex micropollutants in the fjord.
- Antibiotics and fungicides significantly shape bacterial and fungal communities.
- Wastewater impacts reduce microbial diversity and disrupt functional potential.
- Marine sites show microbial recovery and enrichment away from discharge points.
- eDNA and toxic unit modeling link chemical stress to microbiome restructuring.

**Graphical abstract:** 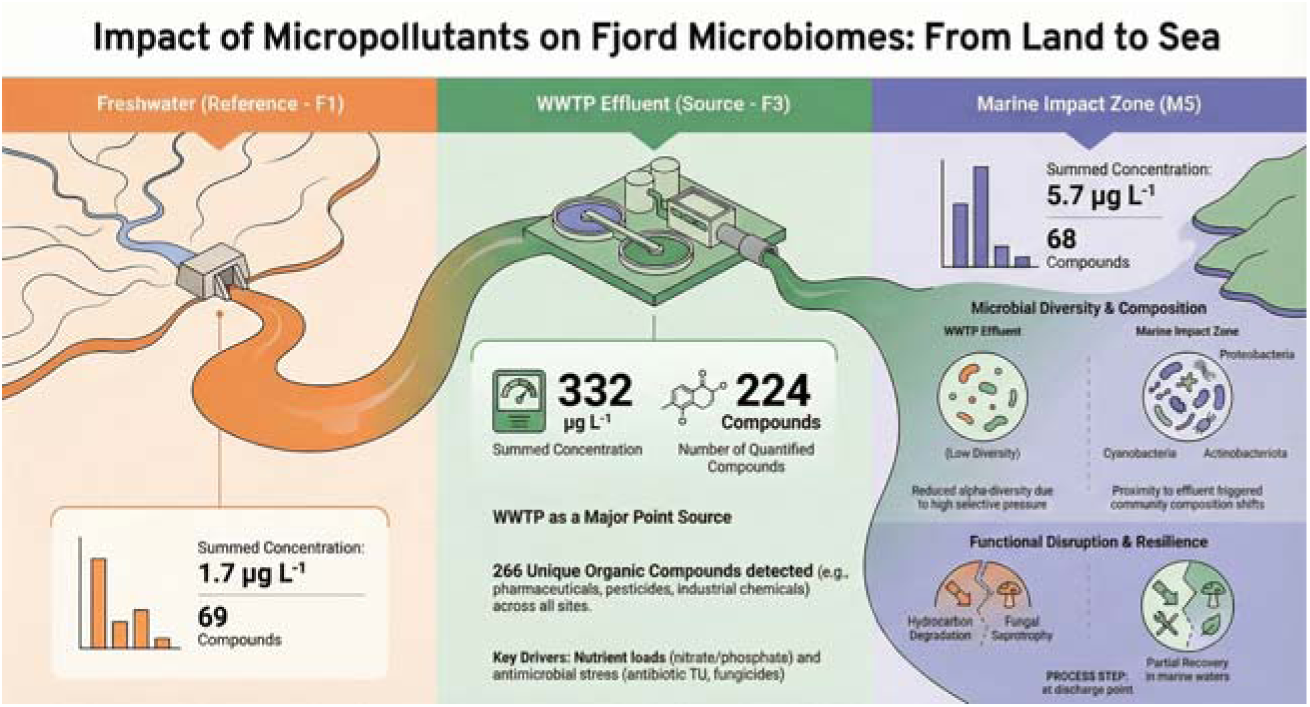

## 1. Introduction

Anthropogenic activities continuously introduce mixtures of micropollutants, including pesticides, pharmaceuticals, and industrial chemicals, into freshwater and coastal environments. Primary sources of these micropollutants in lotic systems include urban and agricultural runoff (Reiber et al., 2021) and the discharge of untreated or treated sewage from wastewater treatment plants (WWTPs) (Beckers et al., 2018). Furthermore, both nutrients and micropollutants are significantly discharged by WWTPs to streams and rivers (Ademollo et al., 2021; Inostroza et al., 2024), as well as to coastal environments (Inostroza et al., 2023a). Despite the implementation of advanced removal systems in WWTPs, some micropollutants persist and can threaten biodiversity across trophic levels, even at low environmental concentrations (Schwarzenbach et al., 2006; Tarigan et al., 2025).

Microbial communities drive ecological processes and are fundamental to the biogeochemical cycling of carbon, nitrogen, sulphur, and other elements (Hutchins and Capone, 2022; Jackson and Gabric, 2022). Rivers, streams, and coastal areas rely on functional microbial communities with high diversity and multifunctionality, enabling differential community responses to allochthonous matter. For example, bacteria, microeukaryotes, and diatoms respond differentially to environmental disturbances, thereby regulating ecosystem health (Clark et al., 2020; Kelly et al., 2024). Microbial metabolism affects all other organisms in their surrounding environment and provides critical ecosystem services such as organic matter decomposition, nutrient recycling, and the degradation of toxic chemicals.

Bacteria and fungi play key roles in biodegradation processes, enzymatically breaking down complex organic molecules and converting them into simpler, and eventually less toxic, forms. Among bacteria, certain genera inhabiting aquatic environments, including species of *Pseudomonas*, *Geobacillus*, *Bacillus*, *Escherichia*, *Rhodococcus*, and *Exiguobacterium*, can degrade micropollutants and their associated metabolites by utilising them as carbon and nitrogen sources (Narayanan et al., 2022). Similarly, fungi, the most efficient degraders in polluted water, play a key role in the biotransformation of persistent pollutants due to their broad enzymatic repertoire and secretion capacity (Ferrando-Climent et al., 2015). These biological transformation processes contribute to the resilience and stability of aquatic environments (Hilderbrand and Utz, 2015; Rodrigues-Filho et al., 2023). However, certain micropollutants, such as antimicrobials (antibiotics, biocides, and bactericides), are specifically designed to disrupt microbial biological functions. In the case of antibiotics, these compounds can drive the development of resistance in the environment (Corno et al., 2019). Owing to their persistence, the non-target effects of these bioactive substances on microbial communities remain a pressing ecological concern (Carusso et al., 2018).

The effects of micropollutants on bacterial and fungal communities are diverse, and various studies report distinct alpha- and beta-diversity patterns (within- and between-sample diversity), as well as shifts in community composition along pollution gradients. In streams and rivers, low bacterial alpha-diversity has been observed at effluents with high micropollutant loads, suggesting a negative impact from urban and agricultural areas. At the community level, variations in bacterial beta-diversity were better explained by nutrient availability and antibiotic stress (Inostroza et al., 2025; Verbücheln et al., 2025). Conversely, high bacterial alpha-diversities have been measured downstream of WWTPs, suggesting differential effects on microbial biodiversity among contaminated effluents (Burdon et al., 2020; Mansfeldt et al., 2020). Similarly, fungal alpha-diversity has been reported to increase downstream of WWTPs (Zhu et al., 2021). However, the community structure of fungi in other WWTP-influenced environments, such as coastal areas, appears distinct from those of bacteria. In these areas, factors such as salinity and nitrogen concentrations have been linked to the abundance and beta-diversity of planktonic fungi (Wang et al., 2019). Despite these insights, micropollutants remain an understudied abiotic factor potentially shaping multi-domain microbiomes in coastal environments receiving human-derived urban effluents. Fjords, characterised by deep, glacially carved valleys with restricted water exchange, are particularly susceptible to micropollutant accumulation and represent natural laboratories for studying pollution gradients in coastal environments (Ademollo et al., 2021; Inostroza et al., 2023a).

Here, we investigated the ecological status of bacteria and fungi in a fjord system (Hakefjorden and Askeröfjorden), including the freshwater inputs from one WWTP and two streams running through agricultural and industrial areas near Stenungsund, located on the Swedish west coast. Specifically, we assessed: (i) structural changes in bacterial and fungal communities due to micropollutant exposure, (ii) the degree to which environmental parameters in the water column shape those structural changes, with a focus on antimicrobial micropollutants, and (iii) potential microbial functional changes associated with micropollutant exposure. We focused on these two fjords because the municipality of Stenungsund hosts the largest chemical cluster in Sweden, producing a wide range of industrial chemicals and representing a major source of potential anthropogenic chemical pressure on adjacent coastal ecosystems. Additionally, this municipality discharges treated sewage effluents into the Hakefjorden fjord system where hundreds of micropollutants, along with legacy chemicals, have been detected in surface waters (Fick et al., 2011; Gustavsson et al., 2017; Inostroza et al., 2023a).

## 2. Material and methods

### 2.1. Sample collection across the Hakefjorden and Askeröfjorden fjord system

Surface water samples were collected at nine sampling sites along the coast of Stenungsund and surrounding localities in October 2020 (fall season), as shown in Figure 1. Marine sites were designated as M1 through M6, progressing northward. M1 was located distant from Stenungsund city, near the Kattegat Sea, while M2 was located near F1, M3 near F2, M4 before the marine WWTP outflow, M5 near the marine WWTP outflow, and M6 further north from M5. Freshwater sites F1, F2, and F3 were selected to represent the influence of agricultural areas, urban and industrial runoff, and treated WWTP effluent, respectively.

A total of 2.5 L of surface water was sampled from two meters below the surface using a Niskin bottle (2.5 L, General Oceanics) per marine sampling site, while 1 L of surface water was sample from freshwater sites. No rain was recorded immediately before, during, or after the sampling campaign. Water was promptly transferred to previously autoclaved glass bottles and stored at 4 °C until reaching the laboratory on the same day. Once in the laboratory, 2 L of water was passed through 0.22 µm Sterivex® filters (Merck, Germany). After filtration, Sterivex filters were immediately stored at −20 °C until subsequent environmental DNA (eDNA) extractions (Chiang and Inostroza, 2022). In addition to the water for eDNA, *in situ* measurements of temperature, conductivity, and pH were recorded using a HANNA HI9811-5 waterproof portable meter. Furthermore, 100 mL of surface water samples were collected for nitrite, nitrate, ammonium, phosphate, silica, and dissolved organic carbon (DOC) measurements at Eurofins Scientific (Sweden).

### 2.2. DNA extraction, sequencing, bioinformatic, amplicon analyses, and functional annotation

The eDNA from water was extracted using the DNeasy PowerWater Kit (Qiagen, Germany), following a slightly modified version of the manufacturer’s protocol to improve DNA yields (Chiang and Inostroza, 2022). The DNA samples were cleaned up and concentrated using AMPure XP clean-up kits (Beckman Coulter, USA) following the manufacturer’s protocol. Bacterial 16S rRNA gene sequences were amplified using the V3-V4 primer pair (341F: CCTAYGGGRBGCASCAG; 806R: GGACTACNNGGGTATCTAAT). The fungal nuclear ribosomal internal transcribed spacer 2 (ITS2) was amplified using the primer pair (ITS3-2024F: GCATCGATGAAGAACGCAGC; ITS4-2409R: TCCTCCGCTTATTGATATGC). The PCR amplification and amplicon sequencing were conducted by Novogene Co. (Cambridge, UK). All samples were sequenced using a 250-bp paired-end Illumina platform.

The raw forward and reverse sequences in “.fastq” format obtained from the 16S rRNA gene (bacteria) and the ITS2 region of the ribosomal operon (fungi) were imported into the software “Quantitative Insights Into Microbial Ecology 2” (QIIME2 2024.5 Amplicon Distribution) (Bolyen et al., 2019). The primer sequences were removed from the libraries using the function “cutadapt trim-paired,” and sequences with an error rate higher than zero were discarded. After trimming, the sequences were dereplicated, denoised, and merged into amplicon sequence variants (ASVs), calling the function “denoise-paired” from the DADA2 plugin (Callahan et al., 2016). This step included chimera filtering of ASVs using the “isBimeraDenovo” method, which detects chimeras based on the consensus chimeric fraction across all samples. Singletons and doubleton were not removed from our datasets. The taxonomic assignment of bacteria was performed with the function “feature-classifier classify-consensus-vsearch” (Rognes et al., 2016), using a cut-off value of 0.8 and 97% identity with the SILVA database version 138.1 (Quast et al., 2012) as reference. The classification of fungi was established through prior training of a classifier using the machine learning algorithm of the function “feature-classifier fit-classifier-naive-bayes” (Pedregosa et al., 2011), and the non-redundant version 10.0 of the UNITE+INSDC fungal ITS database (Abarenkov et al., 2024) as reference.

Among classified sequences, we include the top ten taxa in relative abundance plots; groups below this threshold are mentioned in the results section as “Others”. Furthermore, the ASVs classified as Eukaryota, Metazoa, Archaeplastida, Fungi, and unassigned were excluded from further analyses of the bacterial dataset. Similarly, the ASVs classified as Alveolata, Metazoa, Protista, Viridiplantae, Stramenopila, Eukaryota_kgd_Incertae_sedis, Rhizaria, Cryptista, Ichthyosporia, Amoebozoa, Choanoflagellozoa, Haptista, and unassigned were excluded from further analyses of the fungal dataset. Aiming to confirm the different clustering patterns of both studied communities, the bacterial ASVs were merged with the fungal ASVs to assess both microbial communities in further multivariate analyses using the methods “feature-table merge” and “feature-table merge-seqs”.

Putative ecological function profiles from the 16S rRNA dataset were generated using the FAPROTAX database and its algorithm “collapse table.py” (Louca et al., 2016). The bacterial dataset was analysed, focusing on hydrocarbon degradation (aliphatic non-methane degradation, aromatic compound degradation, aromatic hydrocarbon degradation, and hydrocarbon degradation) and putative energy sources (aerobic chemoheterotrophy, autotrophy, chemoautotrophy, and fermentation profiles). For fungi, the functional analysis of the ITS2 dataset was performed using FUNGuild v1.1, a program designed to assign ecological functions to fungal sequences (Nguyen et al., 2016). For the fungal dataset, the analysis focused on ecological functions related to different energy sources, for example Pathotrophy, Symbiotrophy, Saprotrophy, and diverse types of Saprotrophy.

### 2.3. Assessment of micropollutants

Surface water samples were simultaneously collected for micropollutant characterisation during the same sampling campaign. Sample collection methodology, analytical extractions, and measurements were published in a separate data descriptor manuscript (Inostroza et al., 2023a).

We aimed to assess the pressure of micropollutants on bacterial communities using the well-established toxic unit (TU) approach. However, effect data on bacteria is scarce and limited to antibiotics through the endpoint minimum inhibitory concentration (MIC) of antibiotics (Bengtsson-Palme and Larsson, 2016). Therefore, we normalised environmental antibiotic concentrations by their MIC as suggested by Inostroza et al. (2025) and implemented in Verbücheln et al. (2025). The antibiotic stress (TU_MIC_) was thus calculated as follows:

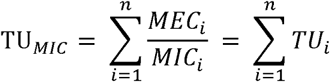

where *MEC*_i_ denotes the measured environmental concentration of antibiotic *i*, while *MIC*_i_ represents the corresponding minimum inhibitory concentration (MIC) of antibiotic *i*. The ratio *MEC*_i_/*MIC*_i_ provides a dimensionless measure of the individual toxicity contribution of each antibiotic compound in the sample. Furthermore, we selected the lowest MIC because antibiotic concentrations below the MIC have been shown to select for resistant bacteria (Andersson and Hughes, 2012; Gullberg et al., 2014). Gullberg et al. (2011) reported that minimal selective concentrations (MSC), defined as the lowest antibiotic concentration at which resistant bacterial strains gain a competitive advantage, ranged from 1/230 to 1/4 of the corresponding MIC, depending on the antibiotic. This highlights that selective pressures can occur at concentrations far below inhibitory levels and vary widely among antibiotics.

Fungi are the intended targets of fungicides. However, comprehensive ecotoxicological data for this group remain limited, posing challenges for environmental risk assessment. Due to this data scarcity, we adopted a pragmatic approach by summing the environmental concentrations of all detected fungicides to generate a fungicide vector hereafter referred to as “summed fungicides”. This vector, representing the total measured fungicide load at each sampling site, was used as a proxy for overall fungicidal stress in subsequent canonical correspondence analyses (CCAs). This approach enabled the exploration of relationships between fungicide and antibiotic occurrence and microbial community composition, analogous to the methodology employed by Verbücheln et al. (2025).

### 2.4. Statistics and reproducibility

The alpha and beta diversity were determined by importing the representative sequences, frequency table, assigned taxonomy, phylogenetic tree, and metadata table of the ASVs into R ver. 4.3.0 (R Core Team, 2024) using the “file2meco” package and processed using the “microeco” package (Liu et al., 2021). We compared the dissimilarity between the microbial communities at each sampling site based on their occurrence. For this, we used the UniFrac unweighted distances in principal coordinate analysis (PCoA), including both bacterial and fungal communities as well as each microbial community individually. Furthermore, canonical correspondence analysis (CCA), a constrained ordination method, was used to evaluate the association between the environmental parameters measured at each station (independent variables) and the bacterial/fungal taxa (dependent variables) detected through metabarcoding. To reduce multicollinearity among explanatory variables prior to CCA, we calculated the variance inflation factor (VIF) and excluded variables with VIF values >1.79, following a conservative threshold to improve model interpretability. Following CCA, we performed a permutation-based analysis of variance (ANOVA) with 999 permutations to assess the statistical significance of the constrained model, including the influence of environmental variables, sampling site differences, and taxa composition.

To further summarize patterns in chemical exposure, we performed a principal component analysis (PCA) on the concentration matrix of detected micropollutants. Prior to ordination, compounds detected in fewer than six of the nine sampling sites were excluded to remove rare or site-specific outliers, and the remaining concentration data were Hellinger-transformed to standardize the scale and reduce the influence of dominant compounds. The first principal component (PC1), representing the major gradient in micropollutant composition across sites, was subsequenctly included as an explanatory variable in additional CCA models for bacterial communities, fungal communities, and both communities combined, to support and validate the main CCA results.

The datasets for our high-throughput sequencing libraries (raw reads) were deposited in the Sequence Read Archive (SRA) in the National Library of Medicine (NCBI) under the accession numbers PRJNA1236954 (16S rRNA) and PRJNA1236969 (ITS2 rRNA).

## 3. Results and discussion

### 3.1. Micropollutant characterisation of the Hakefjorden and Askeröfjorden fjord system

Detailed tables of detected and quantified micropollutants, including their concentrations, and chemical identifiers, are provided in a separate data descriptor (Inostroza et al., 2023a) and are publicly available through the Zenodo repository (Inostroza et al., 2023b).

Micropollutants co-occurred across all freshwater and marine sites, with the highest concentrations observed at the WWTP effluent site F3, indicating the WWTP as a major source of chemical inputs to the Hakefjorden fjord system (Figure 2). Total summed concentrations increased along the freshwater gradient from F1 (1.7 µg L^−1^) to F2 (170 µg L^−1^), reaching a maximum at F3 (332 µg L^−1^) (Figure 2A). Across marine sites, summed micropollutant concentrations were highest at M2 (82.4 µg L^−1^), while most other sites showed comparatively low and similar concentrations (1-2 µg L^−1^), except for M5 (5.7 µg L^−1^). Of the 861 organic compounds analysed, 266, including pharmaceuticals, pesticides, and industrial chemicals, were detected above their method detection limits at least at one site. The number of quantified chemicals varied across stations; freshwater sites ranged from 69 compounds at F1 to 224 at F3 (WWTP effluent), with 114 detected at F2 (Figure 2A). Among marine sites, between 52 compounds (M6) and 68 (M5, the marine site closest to the WWTP effluent F3) were quantified.

The top 20% single-compound concentrations are shown in Figure 2B. These micropollutants were dominated by industrial chemicals, pharmaceuticals, and food- and beverage-related compounds, indicating multiple and diverse pollution sources. The highest single-compound concentrations were observed for 5-methyl-1H-benzotriazole (5-MBT), an industrial chemical, reaching 150 µg L^−1^ at site F2 and 134 µg L^−1^ at the WWTP effluent site F3. Other high-concentration micropollutants included: pharmaceuticals (e.g., metoprolol, norfloxacin, fluconazole, dydrogesterone, and gabapentin), pesticides (e.g., quinmerac, chloridazon, and terbuthylazine-2-hydroxy), and food- and beverage-related compounds (e.g., sucralose and cotinine).

Micropollutants are increasingly detected in coastal environments due to discharges from WWTPs, as well as inputs from rivers and streams. The micropollution pattern observed in the Hakefjorden and Askeröfjorden fjord system, characterised by the widespread co-occurrence of diverse micropollutant classes, is consistent with patterns reported for other fjords and coastal regions (Balakrishna et al., 2023; Choi et al., 2020; Inostroza et al., 2024; Nödler et al., 2014). High concentrations of benzotriazoles, a class of industrial anticorrosive agents, and particularly 5-methyl-1H-benzotriazole (5-MBT), have also been reported in coastal environments (Balakrishna et al., 2023; Yao et al., 2022a). In the Hakefjorden and Askeröfjorden system, the elevated levels of 5-MBT and 1H-benzotriazole likely reflect regional industrial activity in this region given that Stenungsund represents the largest petrochemical hub in Scandinavia and these compounds are widely used as a metal anticorrosive agents and ultraviolet stabilisers in industrial and commercial applications (Cantwell et al., 2015). The co-occurrence and elevated concentrations of micropollutants at freshwater sites F2 and F3 further support the contribution of urban discharge and WWTP effluents as major sources of chemical inputs into the system (Beckers et al., 2020; Soriano et al., 2024; Vidal-Dorsch et al., 2012).

### 3.2. Bacterial and fungal community composition

A total of 311,160 (bacteria) and 832,578 (fungi) sequences were obtained, reaching saturation in rarefaction curves at all sampling sites and indicating sufficient sequencing depth (Figure S1). For bacterial communities, comprised 3,489 and 3,072 amplicon sequence variants (ASVs) in freshwater and marine sites, respectively. For fungi, sequencing yielded 4,218 and 1,657 ASVs in freshwater and marine sites, respectively. Taxonomic assignment identified 2,300 bacterial ASVs and 370 fungal ASVs in marine sites, with the lowest ASV richness observed in the WWTP effluent site F3. Overall, 1,706 bacterial ASVs and 3,276 fungal ASVs remained unclassified across the respective datasets. Detailed information on ASVs and sequence counts per site is provided in Table S2.

Despite differences in taxonomic resolution between operational taxonomic units (OTUs) and ASVs, we compared our results with existing studies using both metrics, as such approaches are commonly used to approximate patterns of biological diversity (Kerrigan and D’Hondt, 2022). Comparable diversity levels have been reported for bacterial communities in WWTP effluent-affected streams and rivers, where more than 300 and up to 760 OTUs per sample were detected upstream and downstream of WWTP effluents, respectively (Zhu et al., 2021). In marine microbiomes, non-polluted coastal areas have been shown to harbor higher bacterial ASV richness than observed in our study (He et al., 2023). However, bacterial ASV richness can increase markedly in areas characterised by elevated micropollutant concentrations, consistent with the higher ASV richness observed at site M5 compared to other marine sites (Table S2). In contrast, fungal ASVs richness was reduced at site F3 relative to F1 and F2, whereas site M5 exhibited notably higher fungal ASV richness than other marine sites (Table S2). Similarly, reduced fungal genotype richness (i.e., ASVs or OTUs numbers) has been reported in WWTP-affected streams impacted by micropollutants (Li et al., 2023).

Bacterial and fungal community composition revealed that, at the phylum level, the WWTP effluent site F3 was markedly distinct from F1 and F2 (Figure 3). These differences were consistent across deeper taxonomic levels, including order and family, for both microbial groups (Figure S2 and S3). From a bacterial perspective, the phylum Bacteroidota–one of the most prevalent bacterial groups alongside Proteobacteria–showed reduced relative abundances at F3 (16%), accompanied by increased relative abundances of Campylobacterota (14%), Patescibacteria (9%), and Firmicutes (7%) (Figure 3A). Similar shifts involving Bacteroidota, Firmicutes, and Campylobacterota have been reported in streams receiving treated wastewater effluents (Mansfeldt et al., 2020).

Across marine sites (M1-M6), bacterial communities exhibited a more stable structure than those observed in freshwater sites. Notable changes occurred at site M5, the location closest to the WWTP outflow, where Cyanobacteria and Actinobacteriota showed reduced relative abundances (6% and 4%, respectively) compared to other marine sites (8-15%). This reduction coincided with an increase in Proteobacteria which reached up to 53% relative abundance. Similar increases in Proteobacteria have been documented in coastal waters receiving WWTP effluents in the Baltic Sea (Vaquer-Sunyer et al., 2016).

At the order level, bacterial community composition among freshwater sites was characterised by a strong dominance of the category “Others” (>70%) at site F3 (i.e., taxa outside to the ten most abundant orders), accompanied by a moderate increase in Microtrichales (3%) and a pronounced decrease in Flavobacteriales (5%) compared to F1 and F2 (>20%; Figure S2A). A similar pattern was observed at the family level, with an increased contribution of “Others” (>75%), a marked reduction in Flavobacteriaceae (5%), and the complete absence of Spirosomaceae (Cytophagales) and Methylophilaceae (Methylopilales) at the WWTP effluent site F3 (Figure S2B).

Consistent with these observations, Mansfeldt et al. (2020), reported that bacterial groups belonging to Bacteroidota (e.g., Flavobacteriales and Spirosomaceae) are scarce in WWTP effluents. In contrast, other studies have documented increased abundances of these groups under similar conditions, particularly Flavobacterales, which are often dominant in WWTP and are involved in the degradation of complex organic matter such as cellulose (Burdon et al., 2020; Kim and Yu, 2020). Our results suggest that WWTP effluents can exhibit distinct chemical fingerprints, which may differentially shape the composition of these bacterial groups.

At marine sites, changes at the order level were subtle and included a sligh increase in the category “Others” group (>25%) and a concomitant decrease in Synechococcales (5%) at site M5, the location closest to the WWTP effluent (Figure S2A). A similar pattern was observed at the family level, with an increased contribution of “Others” (>40%) and a reduced relative abundance of Cyanobiaceae (5%) (Synechococcales, Cyanobacteria) at M5 (Figure S2B). Members of Synechococcales, a major phytoplankton lineage, are ubiquitous across marine environments and are known to vary along coastal gradients primarily in response to temperature (Doré et al., 2022). However, in coastal systems influenced by WWTP effluents, this cyanobacterial group has also been reported to positively correlate with nutrient loads (Vaquer-Sunyer et al., 2016). These observations suggest that local physicochemical conditions associated with effluent proximity may modulate the response of Synechococcales in coastal microbial communities.

For fungi, the dominant phylum Basidiomycota showed a pronounced decrease in relative abundance at site F3 (2%), whereas Chytridiomycota (10%) and the category “Others” (> 20%) increased (Figure 3B). Although information on fungal community composition in WWTP effluents remains limited, Basidiomycota has been consistently reported as a prevalent fungal group in freshwater streams (Li et al., 2023).

Across marine sites, fungal communities displayed site-specific patterns, particularly at M1 and M5. At M1, an increased contribution of the “Other” fungal category (15%) was observed, whereas at M5, relative abundances of Chytridiomycota (8%) and Rozellomycota (5%) were elevated (Figure 3B). The increase in Chytridiomycota at M5, potentially associated with the proximity to the WWTP outflow at F3, may reflect the dominance of this fungal group in WWTPs, where Chytridiomycota–known for their capacity to degrade chitin and keratin–can account for up to 40% of total fungal relative abundance (Assress et al., 2019).

At the order and family levels, fungal communities at marine sites show a clearly distinct community composition compared to freshwater sites (Figure S3). In the WWTP effluent site F3, fungal orders were almost entirely dominated by the category “Others”, accompanied by marked decrease in Hypocreales and Leotiales, both of which were dominant at sites F1 (12%) and F2 (7%). In contrast, the relative abundance of Saccharomycetales at F3 was comparable to that observed at F1 and F2 (up to 0.5%) (Figure S3A). Previous studies have reported that fungal orders such as Hypocreales are among the dominant taxa in wastewater treatment plants (Buratti et al., 2022; Korniłłowicz-Kowalska et al., 2022).

Among marine sites, fungal orders such as Rhizophydiales (Chytridiomycota) were primarily detected at M3 (18%), M4 (5%) and M5 (11%), distinguishing these three sites from the remaining marine locations. (Figure S3B). In addition, the orders Pleosporales, Polyporales, Agaricales, Helotiales, Hypocreales, Russulales, Saccharomycetales and Eurotiales collectively distinguished site M5 from all other marine sites. Taken together, these patterns suggest that proximity to the WWTP effluent may influence the composition of fungal communities at M5, including taxa recognized for their roles in the degradation of complex organic compounds in the environment (Cortés-Lorenzo et al., 2016; Eichlerová and Baldrian, 2020; Usharani, 2019; Zhang et al., 2022).

At the family level, fungal communities at all freshwater sites were dominated by the category “Others”, particularly at site F3. Sites F1 and F2 exhibited similarly low relative abundances of Typhulaceae, Coralloidiomycetaceae, Aspergillaceae, Herpotrichellaceae, and Phaeosphariaceae (>1%) whereas these families were further reduced at F3 (<1%). Among marine sites, site M5 differed markedly from the remaining locations, showing reduced relative abundances of families that were otherwise dominant in marine environments, such as Agaricaceae (6%) and Pezizellaceae (6%). In contrast, M5 was characterised by higher relative abundances of Typhulaceae (14%) and the “Others” category (60%) (Figure S3B). Although information on these fungal families in marine systems remains limited, the detection of Agaricaceae is noteworthy, as this family includes saprobic taxa tipically associated with grassland and woodland habitats (Kumar et al., 2021). Taken together, our metabarcoding results reveal previously underreported freshwater and marine fungal assemblages that appear highly sensitive to local environmental conditions, potentially influenced by WWTP effluent inputs at site M5.

### 3.3. Bacterial and fungal diversities in fjords impacted by micropollutants

Our results revealed distinct patterns in bacterial and fungal alpha-diversity that differentiated freshwater from marine sites, with notable high alpha-diversity values observed at sites F3 and M5, respectively (Figure 4). Overall, bacterial alpha diversity was higher in freshwater than in marine sites, except for M5 (Shannon index = 7.8; Figure 4A), a marine site influenced by WWTP outflow that exhibited elevated richness relative to the other marine locations.

Among freshwater sites, F1 was identified as the least affected by micropollutants and accordingly exhibited the highest ASVs richness (F1: 1,358 ASVs) compared to F2 (1,074 ASVs) and the WWTP effluent site F3 (1,057). In contrast, F3 displayed the highest Shannon diversity (F3: 8.6 vs. F1: 8.5 and F2: 8.4; Figure 4A) and evenness (F3: 0.855 vs. F1: 0.820 and F2: 0.825), indicating a more even distribution of taxa despite lower richness. Similar patterns have been reported for riverine ecosystems impacted by urban and agricultural activities, where bacterial richness and diversity are reduced under chemical stress (Inostroza et al., (2025), in agreement with our observations. In contrast to bacteria, fungal communities exhibited the highest alpha-diversity values at freshwater sites F1 and F2, whereas site F3 displayed markedly lower diversity, comparable to that observed at marine sites (Figure 4B). Specifically, observed ASV richness reached up to 1,800 at F2 but declined to 280 at F3, while Shannon diversity values approached 9 at F1 and F2 and decreased to 4.2 at F3.

Across marine sites, fungal communities showed relatively similar ASVs richness and Chao1 estimates. However, site M5 exhibited the highest fungal diversity (Shannon index = 6.8 vs. ≤ 4.6 at other marine sites) and evenness (M5: 0.83 vs. ≤ 0.745 at M1-M6). Collectively, these patterns suggest that WWTP effluent at F3 may substantially alter fungal community structure in receiving waters, consistent with previous observations in freshwater streams (Zhu et al., 2021) and in systems exposed to wastewater containing high concentrations of fungicides (Burdon et al., 2020).

Community dissimilarity (beta diversity) was assessed using two complementary approaches: (i) analysing bacterial and fungal communities independently, and (ii) analysing both communities jointly (Figure 5). This strategy allowed us to capture patterns specific to each microbial group while also providing an integrated view of community structure. When analysed separately, both bacterial and fungal communities formed three well-defined clusters (Figures 5A and 5B). One cluster was dominated by marine sites, whereas another was dominated by the freshwater sites F1 and F2 (agricultural and industrial/urban, respectively). Notably, the WWTP effluent site F3 formed a distinct cluster for bacteria (Figure 5A), but clustered closely with the site M5 in the fungal community ordination (Figure 5B). When bacterial and fungal communities were combined, the two groups separated clearly in multivariate space, with no overlap between bacterial and fungal assemblages (Figure 5C). Within each group, freshwater and marine sites remained distinct, with F3 and M5 clustering together for bacteria. This combined analysis confirmed the taxonomic separation of the two target groups and provided an alternative perspective on community structuring across environments. Similar patterns of community dissimilarity along environmental gradients, particularly under biocides and wastewater-related stress, have been reported for bacterial and fungal communities in WWTP-affected systems (Burdon et al., 2020; Inostroza et al., 2025; Yao et al., 2022).

Analysis of ASVs shared between freshwater and marine sites revealed two main patterns: (i) elevated number of shared ASVs at the receiving marine site M5 for both bacterial and fungal communities, and (ii) a high degree of ASVs sharing among marine sites (Figure S4). Overall, only moderate numbers of ASVs were shared between freshwater plume sites (F1, F2, and F3) and receiving marine sites, with even fewer shared ASVs detected at marine sites located upstream of the freshwater plumes, for both bacteria and fungi. These results partially contrast with observations from riverine systems, where ASVs originating from WWTP effluents have been reported to persist downstream in receiving waters (Campbell et al., 2015). In fjord systems, however, enhanced dilution, salinity gradients, and hydrodynamic mixing may limit the persistence and detectability of freshwater-derived ASVs in marine environments, potentially explaining the observed discrepancy.

### 3.4. Association of the bacterial-fungal communities with the environmental parameters

Significant correlations were observed between bacterial-fungal community dissimilarities and multiple environmental parameters (Figure 6). The best-supported model, incorporating nutrients (phosphate, nitrite, nitrate and silica), micropollutants stress (TU antibiotics and summed fungicides), as well as conductivity and pH, explained up to 87.3% of the variability in combined bacterial-fungal community structure. Among the environmental variables, conductivity (R² = 0.6608, p = 0.001), pH (R² = 0.6814, p = 0.001), nitrite (R² = 0.7964, p = 0.041), nitrate (R² = 0.7563, p = 0.022), silica (R² = 0.6439, p = 0.002), phosphate (R² = 0.7620, p = 0.002), and summed fungicides (R² = 0.7817, p = 0.049) were all significantly associated with microbial community variation. Conductivity and pH were positively associated with marine bacterial and fungal communities.

To further summarise micropollution gradients, the first principal component (PC1) derived from the PCA of selected micropollutant concentrations was included in additional CCA models (Figure S6, S7 and S8). Although PC1 was significant, it primarily captured the broad separation between fungal and bacterial assemblages, being positively associated with fungal communities and negatively associated with bacterial communities, except for bacterial communities at the WWTP effluent. In contrast, the main CCA (Figure 6) resolved more specific ecological relationships, showing a positive association between antibiotic stress and bacterial communities at the WWTP effluent, and between fungicides and nitrate with fungal communities at effluent-impacted sites.

Two peaks of antibiotic toxic units were identified within the Hakefjorden and Askeröfjorden fjord system (Table S1), occurring at marine site M2 (TU_MIC_ = 2.2) and at the WWTP effluent site F3 (TU_MIC_ = 1.4), indicating a potential impact of measured antibiotics on microbial communities. Similarly, summed fungicide concentrations peaked at M2 (summed fungicides = 12.28) and were markedly elevated at F3 (summed fungicides = 178.4) (Table S1). Collectively, TU antibiotics and summed fungicides, together with nitrite, nitrate and phosphate, were strongly associated with bacterial and fungal community composition at site F3, whereas silica concentrations were more closely associated with microbial communities at sites F1 and F2.

When bacterial and fungal communities were analysed independently, differences in community structure were primarily explained by pH and conductivity, which were significant drivers of community dissimilarity at marine sites M2, M4, M5 and M6 (Figure S5A). For bacterial communities, TU antibiotics significantly explained community distances at sites M2 and M3, whereas nitrite, phosphate, and silica were the main factors associated with the bacterial community dissimilarity observed at the WWTP effluent site F3. Additional variables associated with bacterial community structure at F3 included summed fungicides, phosphate, and silica. For fungal communities, phosphate and nitrite were significantly associated with community dissimilarity at site F3 (Figure S5B). Other parameters, including nitrate, summed fungicides, and TU antibiotics, were also associated with fungal community dissimilarity at F3, although these relationships were not statistically significant. Conductivity was associated with fungal community dissimilarity across several marine sites (M1, M2, M3, M4 and M6), while pH was primarily associated with fungal community dissimilarity at site M5 and, to a lesser extent, at F2. Silica was the only variable associated with fungal community dissimilarity at site F1, although this association was not significant.

The strong association between conductivity and marine bacterial and fungal communities likely reflects salinity-driven turnover between freshwater and marine habitats, consistent with previous studies documenting salinity as a major determinant of microbial community composition (Liu, 2022; Unno et al., 2015). Phosphorus, a key limiting nutrient in aquatic ecosystems, is primarily cycled by bacteria and has been shown to influence bacterial diversity and community structure at elevated concentrations (LeBrun et al., 2018). This may partly explain why phosphate emerged as a significant environmental factor associated with microbial community variation in our study. Similarly, phosphorous availability has been reported to shape fungal ecology by selecting for specific taxa (Fernandes et al., 2009; Pietryczuk et al., 2018). Nitrogen compounds such as nitrite and nitrate, which are closely linked to microbial metabolic processes, were highest at site F3 (WWTP effluent) and were significantly associated with community dissimilarity. Elevated concentrations at this site may reflect enhanced microbial turnover and nutrient loading typical of wastewater-impacted systems, as reported for bacterial and fungal communities in other freshwater ecosystems (Lear et al., 2014; Miura and Urabe, 2015). Consistent with prior work indicating that bacteria are generally more sensitive to pH variation than fungi (Su et al., 2021), pH was strongly associated with bacterial community structuring in our analyses (Figures S5A and S5B). Summed fungicides were identified as a significant factor associated with overall microbial community composition when bacterial and fungal datasets were analysed jointly. In separate analyses, fungicides were primarily associated with bacterial communities at the WWTP effluent site F3. Previous studies have also reported shifts in freshwater microbial communities in response to fungicide exposure (Burdon et al., 2020), supporting the patterns observed here. Future studies should aim to determine environmental concentration thresholds of fungicides that may significantly influence fungal community structure in freshwater and marine environments. Silica was associated with microbial community variation at several freshwater sites. While anthropogenic inputs may increase silica loads locally (Li, 2017), silica also represent a naturally occurring component of aquatic systems and may act as an indirect indicator of broader geochemical or hydrological gradients rather than a direct stressor.

In the CCAs, several bacterial ASVs were significantly associated with environmental variation across sampling sites. These members of Syntrophobacteria, a group of syntrophic bacteria involved in fermentation and propionate degradation (Westerholm et al., 2022), and Syntrophia, which comprises taxa capable of anaerobic oxidation of aromatic compounds (Langwig et al., 2022). Turneriella (Spirochaetota), known to utilize long-chain fatty acids and fatty alcohols as energy sources (Zuerner, 2015), and Truepera, an extremophilic genus resistant to radiation and metabolically versatile in assimilating sugars, organic acids and amino acids (Albuquerque et al., 2005), were also identified. In addition, ASVs affiliated with Marinimicrobia, a widespread but still poorly characterized marine lineage (Tarn et al., 2016), and Chloroflexi (e.g., strains TK10 and KD4-96), which are broadly distributed in aquatic systems and contribute to carbon cycling through degradation of complex organic compounds (Freches and Fradinho, 2024), were significantly associated with environmental gradients.

Further, Latescibacterota, previously linked to the degradation of persistent organic pollutants (Oliveira et al., 2023), and Planctomycetota (Pla4), reported from environments characterized by antimicrobial resistance hotspots (Godinho et al., 2024), were detected among the environmentally associated ASVs. Collectively, these taxa suggest functional traits related to anaerobic metabolism, organic matter degradation, and potential responses to chemical stressors. Although WWTP effluents are recognized as sources of microbial contaminants and potential pathogens in aquatic environments (Xiao et al., 2024), our data do not directly demonstrate pathogenicity but rather indicate the presence of taxa that may be enriched under wastewater-influenced conditions.

ASVs significantly associated with environmental parameters in the bacterial community analysis are summarised in Table S3. Based on the known ecology of the associated bacterial and fungal taxa, we tentatively inferred potential source signatures, including agricultural inputs, suggested by the presence wood- and hydrocarbon-degrading taxa as well as plant-associated microorganisms– and urban or hospital effluents, indicated by taxa previously reported as opportunistic or potential pathogens (Austin and Moss, 1986; Coorevits et al., 2011; Fei et al., 2015; Fischer-Romero et al., 1996; Fortina et al., 2001; Lee, 2007; Letcher et al., 2006; Nakase et al., 2006; Pozzi et al., 2018; Proia et al., 2004; Ueki et al., 2014; Whitehead et al., 2005). However, inferring environmental sources solely from taxonomic affiliation has inherent limitations, as many microbial taxa are ecologically versatile and may occur across multiple habitats. Moreover, the eDNA metabarcoding approach employed here does not distinguish between active, dormant, or dead organisms. Consequently, the detected ASVs do not necessarily represent metabolically active members of the microbial communities.

### Putative functions in the detected bacterial and fungal genotypes

Bacterial taxa with potential to biodegrade organic chemicals were more abundant at freshwater sites than at marine sites (Figure 7A). High relative abundances of aromatic- and hydrocarbon-degrading functions were observed at F3, F2, and F1, whereas these functional groups were only weakly represented at marine sites M1, M2, M3, and M5. This contrast highlights differences in the ecological functions potentially provided by bacterial communities in freshwater versus marine ecosystems (Cabello-Yeves and Rodriguez-Valera, 2019). Similar enrichment of hydrocarbon-degrading bacteria has been reported in polluted urban river systems (Inostroza et al., 2025), consistent with our observations. In addition, functions related to aromatic hydrocarbon degradation and aliphatic non-methane hydrocarbon degradation were predominantly detected at sites F2 and F3, and to a lesser extent at marine sites M1, M2, and M3.

Regarding bacterial metabolic potential aerobic chemoheterotrophy showed a clear pattern, with higher relative abundances at freshwater sites than at marine sites. However, at the WWTP effluent site F3, the relative abundance of aerobic chemoheterotrophy was reduced to approximately half of the levels observed at F1 and F2. In addition, metabolic functions such as methylotrophy (the utilization of single-carbon compounds) and oxygenic photoautotrophy were detected across all sites, but with notably lower relative abundances at F3 (Figure 7B). Future studies integrating microbial guild interactions (e.g., bacterial-fungal) may help elucidate whether specific classes of micropollutants influence key carbon cycling processes in aquatic ecosystems.

For fungal communities, predicted ecological functions were reduced at the WWTP effluent site F3 compared to other freshwater sites, followed by a relative increase at site M5 compared to the remaining marine locations. The dominant functional guilds included saprotrophy, pathotrophy, symbiotrophy, and wood saprotrophy (Figure 7C). As key decomposers, fungi play a central role in organic matter breakdown and nutrient cycling in aquatic ecosystems. The reduced functional representation observed at F3 suggest a disruption of these processes in wastewater-influenced freshwater environments; however, the partial recovery observed at the nearest marine site (M5) indicates a degree of functional resilience within the fungal community. Similar patterns of increased microbial functional potential under elevated nutrient conditions have been reported for both bacterial and fungal communities (Zhang et al., 2018).

More broadly, our results indicate that wastewater effluents act as strong selective pressures on microbial communities, leading to restructuring of their functional potential. Comparable responses have been documented in WWTP-impacted river sediments, including enhanced anaerobic methane oxidation (Yu et al., 2025). However, the functional responses of fungal communities to WWTP inputs in fjord systems remain poorly documented, and our results provide new insights into these dynamics in coastal environments.

## 3. Conclusion

This study provides a comprehensive assessment of micropollutant contamination and its effects on bacterial and fungal microbiomes in a fjord system. We observed widespread co-occurrence of diverse micropollutants, with the wastewater treatment plant (WWTP) effluent acting as a major point source, characterized by the highest concentrations and diversity of contaminants. This discharge introduces a complex mixture of pharmaceuticals, pesticides, and industrial chemicals, including elevated benzotriazoles associated with regional industrial activities, into freshwater environments and influences adjacent marine microbiomes.

Microbial community analyses revealed pronounced shifts in both bacterial and fungal composition and diversity along the micropollutant gradient. The WWTP effluent harboured distinct microbial assemblages, reflected in altered taxonomic profiles and alpha-diversity patterns. The marine site closest to the effluent plume (M5) exhibited elevated richness and diversity, suggesting selective pressures or enrichment effects associated with wastewater inputs. Beta-diversity analyses further confirmed distinct microbial signatures among freshwater, marine, and WWTP-impacted sites.

Environmental drivers, including nutrients, micropollutants stress (fungicides and antibiotics), and physicochemical factors such as pH and conductivity, were significantly associated with microbial community structure. Taxa indicative of agricultural and urban/hospital sources further highlighted the multiple origin of microbial inputs into the fjord system. Functional predictions suggested that WWTP effluent alters the metabolic potential of both bacterial and fungal communities, reducing key ecological functions at the effluent source while showing partial recovery or functional in receiving marine waters. Collectively, our results demonstrate that WWTP-derived micropollutants mixtures act as strong environmental filters that reshape microbial diversity and functional potential in fjord systems. These findings emphasize the ecological sensitivity of coastal systems to anthropogenic chemical inputs and underscore the need for improved wastewater management and further investigation into the long-term ecological consequences of micropollutant exposure in marine environment.

## 4. Acknowledgements

The processing of raw sequences was enabled by resources in project [SNIC 2022/22-169] provided by the Swedish National Infrastructure for Computing (SNIC) and in project [NAISS 2023/22-5] provided by the National Academic Infrastructure for Supercomputing in Sweden (NAISS) at UPPMAX, funded by the Swedish Research Council through grant agreement no. 2022-06725.

## 5. Author Contributions

CRediT authorship contribution statement: Eduardo Acosta: Methodology, Software, Validation, Formal analysis, Investigation, Data Curation, Writing - Original Draft, Writing - Review & Editing, Visualization. Nicolai Laufer: Software, Writing - Review & Editing. Sonja Schaufelberger: Software, Writing - Review & Editing. Henrik Nilsson: Writing - Review & Editing. Werner Brack: Writing - Review & Editing. Alexis Fonseca: Writing - Review & Editing. Thomas Backhaus: Writing - Review & Editing, Funding acquisition. Pedro A. Inostroza: Conceptualization, Methodology, Validation, Investigation, Data Curation, Writing - Original Draft, Writing - Review & Editing, Supervision, Project administration.

## 6. Declaration of generative AI and AI-assisted technologies in the manuscript preparation process

During the preparation of this work the author(s) used Google Gemini in order to conduct grammar checking and graphical abstract. After using this tool/service, the author(s) reviewed and edited the content as needed and take(s) full responsibility for the content of the published article.

