## Supplementary Material for "Metabarcoding reveals that bacterial and fungal microbiomes are perturbed by micropollutants in a fjord system (Hakefjorden and Askeröfjorden) at the Swedish west coast"

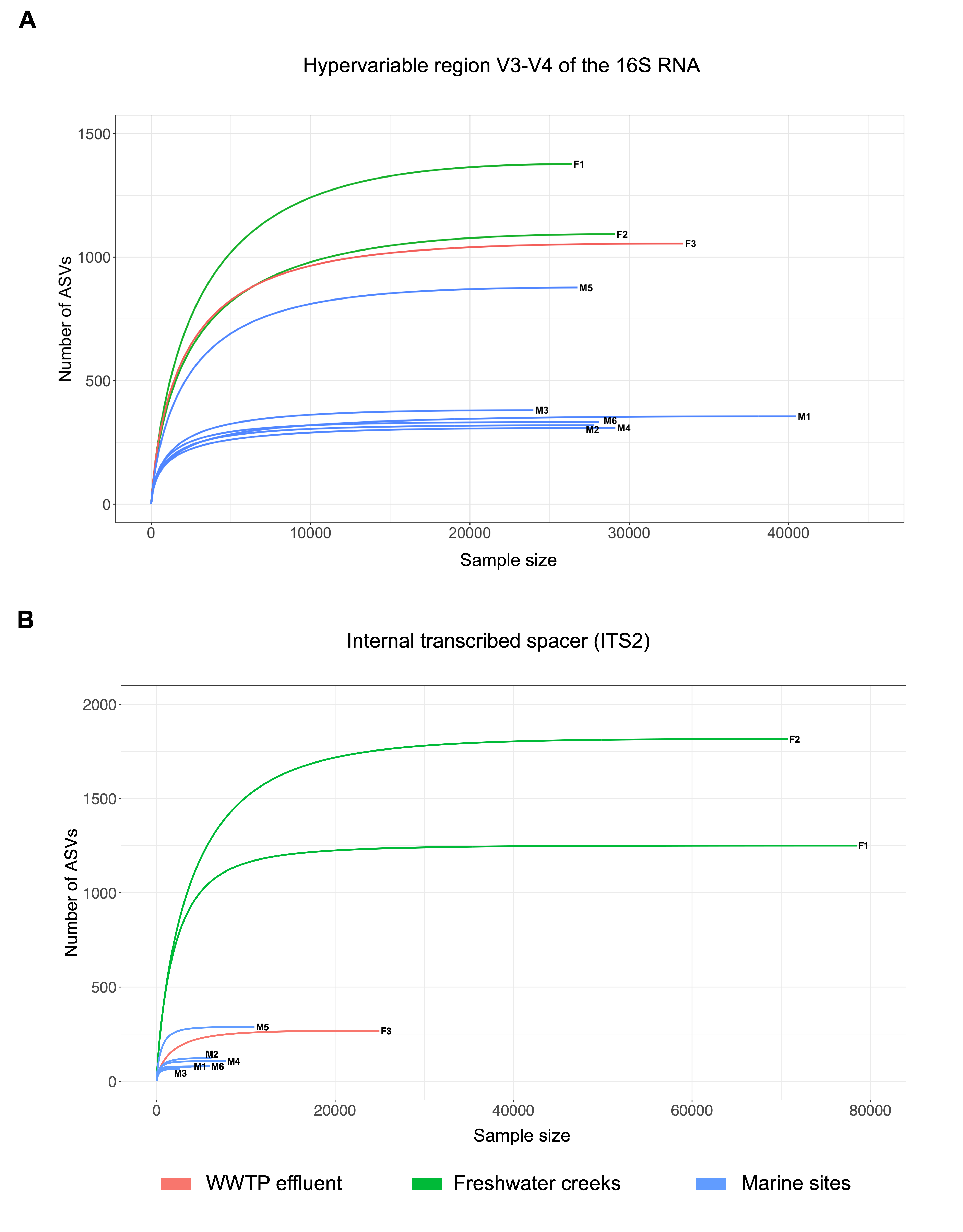

Figure S1. Rarefaction curves. A) Rarefaction curves including the total ASVs of the hypervariable V3-V4 region of the 16S rRNA gene. B) Rarefaction curves including the total ASVs of the internal transcribed spacer 2 (ITS2) of the nuclear ribosomal operon gene. Sampling site types are coloured coded.

Figure S2. Relative abundance of bacterial orders A) and families B) detected in the Hakefjord and Askeröfjord fjord system. Freshwater sites are labelled F1-F3 while marine sites (M1-M6).

Figure S3. Figure S2. Relative abundance of fungal orders A) and families B) detected in the Hakefjord and Askeröfjord fjord system. Freshwater sites are labelled F1-F3 while marine sites (M1-M6).

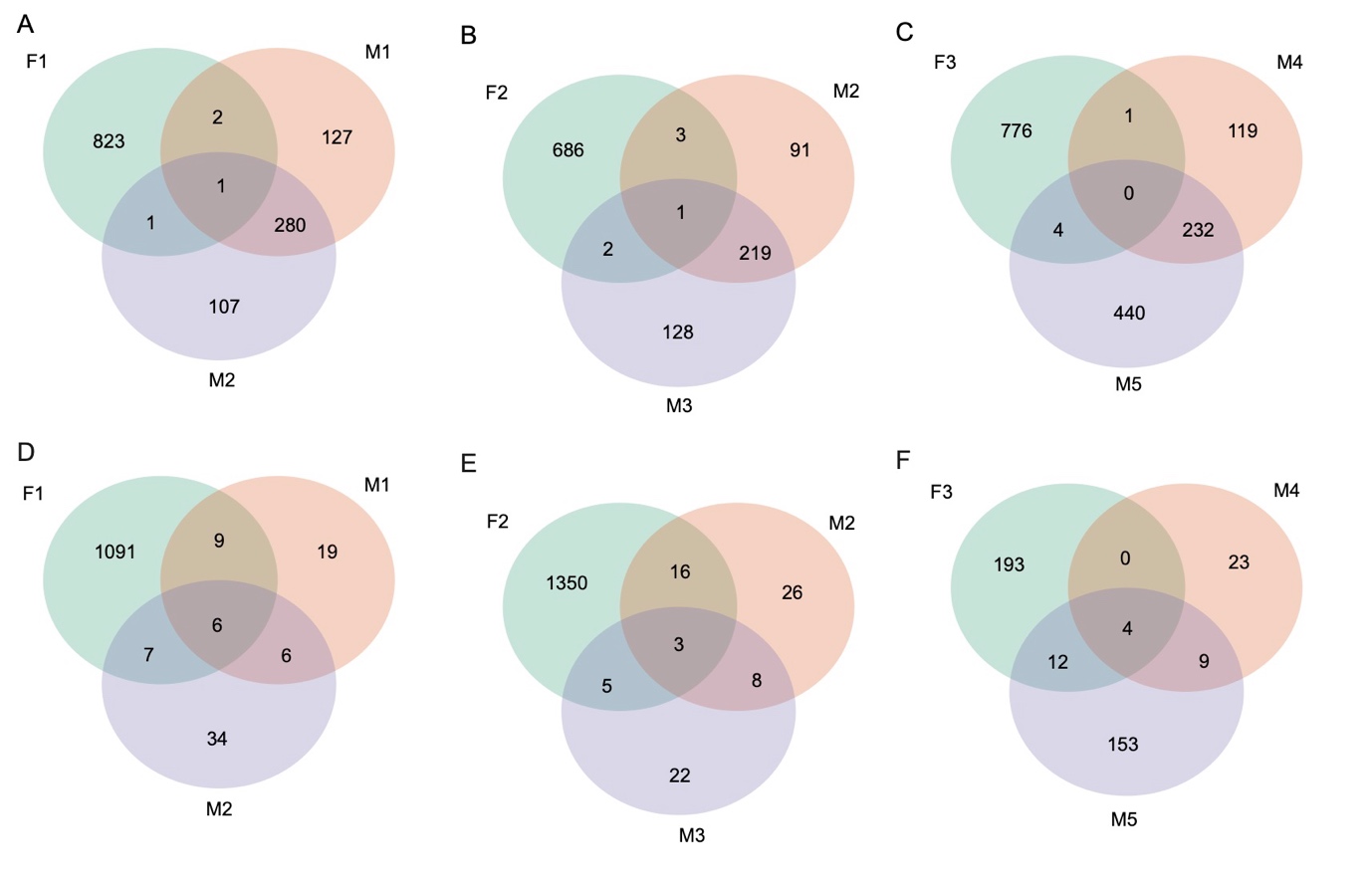

Figure S4. Shared ASVs between freshwater plumes (F1, F2, and F3) and their respective marine sites located before and after the discharges. Shared bacteria in A, B, C and fungi in D, E, and F.

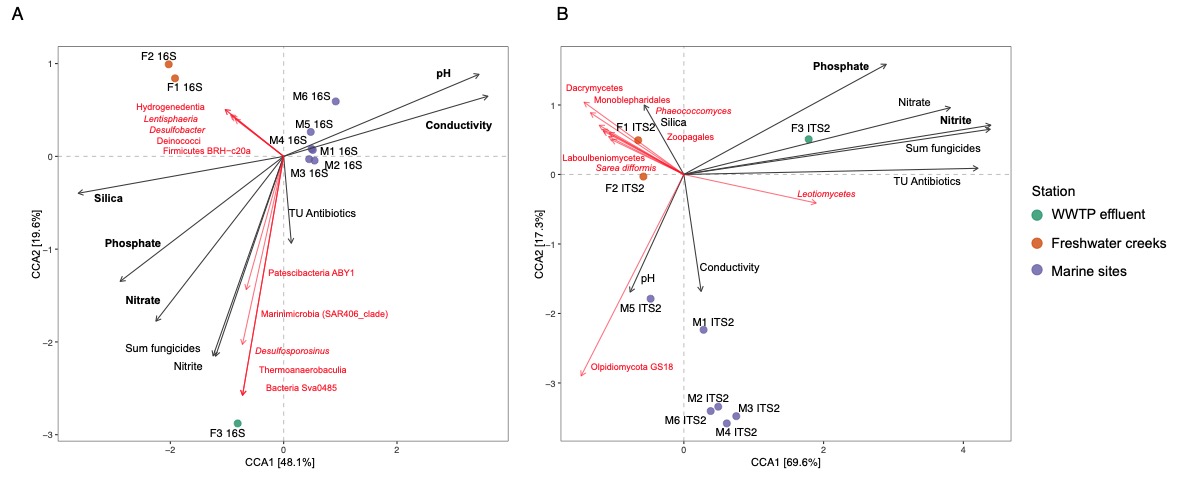

Figure S5. Canonical correspondence analysis (CCA) of the genotypes classified as prokaryotes (A) and fungi (B) detected across stations based on their dissimilarity measured through the UniFrac unweighted distance. The black arrows represent the environmental variables correlated to the studied communities, and the red arrows represent the genotypes associated with the environmental variables.

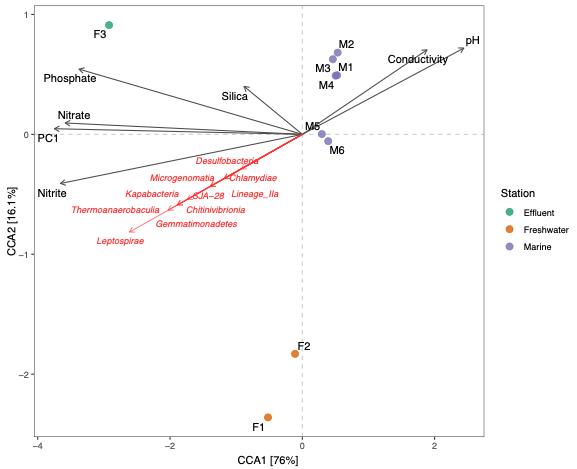

Figure S6. Canonical correspondence analysis (CCA) of the genotypes classified as prokaryotes detected across stations based on their dissimilarity measured through the UniFrac unweighted distance. The variable PC1 derives from the PCA of selected micropollutant concentrations detected in this study.

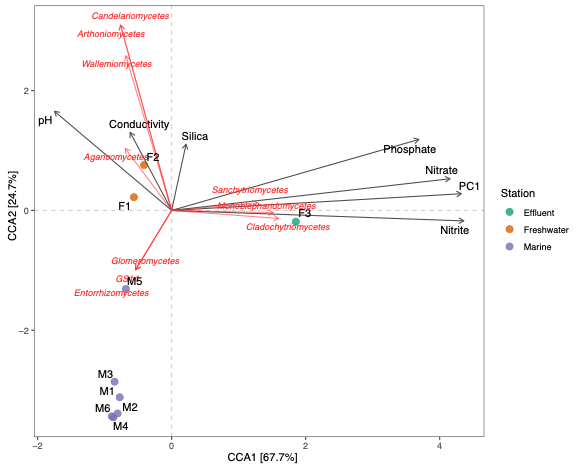

Figure S7. Canonical correspondence analysis (CCA) of the genotypes classified as fungi detected across stations based on their dissimilarity measured through the UniFrac unweighted distance. The variable PC1 derives from the PCA of selected micropollutant concentrations detected in this study.

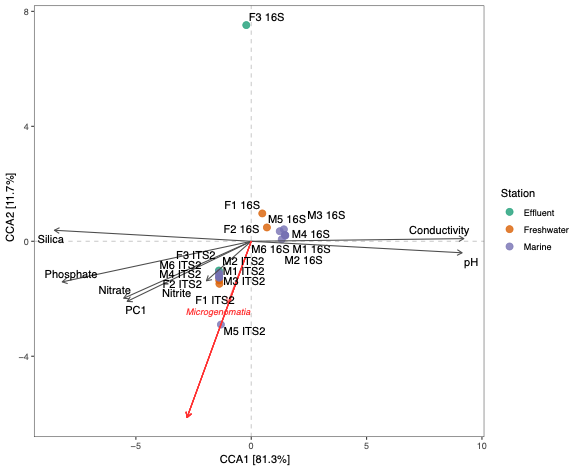

Figure S8. Canonical correspondence analysis (CCA) of the genotypes classified as prokaryotes and fungi together, detected across stations based on their dissimilarity measured through the UniFrac unweighted distance. The variable PC1 derives from the PCA of selected micropollutant concentrations detected in this study.

Table S1. The environmental parameters measured at each sampling site.

| Study Site | M1 | M2 | M3 | M4 | M5 | M6 | F1 | F2 | F3 |
| --- | --- | --- | --- | --- | --- | --- | --- | --- | --- |
| Parameter |  |  |  |  |  |  |  |  |  |
| pH | 8.32 | 8.35 | 8.29 | 8.29 | 8.43 | 8.34 | 6.88 | 6.77 | 6.51 |
| Conductivity (mS/cm) | 33.29 | 34.07 | 33.76 | 34.18 | 36.30 | 35.50 | 0.16 | 0.30 | 0.53 |
| Temperature (°C) | 14.89 | 14.66 | 14.25 | 14.61 | 15.20 | 14.78 | 12.00 | 11.80 | 16.80 |
| Dissolved oxygen (mg/L) | 4.88 | 4.90 | 4.68 | 4.39 | 7.38 | 5.08 | 5.84 | 6.06 | 5.18 |
| Nitrite (µmol/L) | <0.02 | <0.02 | <0.02 | 0.02 | 0.02 | <0.02 | 0.30 | 0.28 | 5.43 |
| Nitrate (µmol/L) | 0.09 | <0.05 | 0.06 | <0.05 | 0.08 | <0.05 | 30.83 | 69.09 | 167.63 |
| Ammonium (µmol/L) | 3.45 | 0.20 | 0.89 | 0.48 | 0.78 | 0.55 | 0.26 | 1.00 | 79.11 |
| Phosphate (µmol/L) | 0.04 | 0.03 | 0.03 | 0.03 | 0.08 | 0.09 | 0.36 | 0.40 | 0.68 |
| Silica (µmol/L) | 0.92 | 1.05 | 1.08 | 1.15 | 3.35 | 2.26 | 100.53 | 150.18 | 107.13 |
| Dissolved organic carbon (mg/L) | 3.82 | 2.66 | <2.50 | <2.50 | 3.46 | <2.50 | 10.30 | 8.33 | 8.39 |
| Antibiotic stress (TU_MIC_) | 0.00925 | 2.20579 | 0.00755 | 0.0068 | 0.0078 | 0.0070 | 0.00004 | 0.0061 | 1.4069 |
| Summed fungicides | 10.9008 | 14.2831 | 4.2 | 3.1 | 2 | 2.3 | 6.66 | 24.94 | 178.4351 |

Table S2. Summary of sequences and ASVs per sampling site before and after taxonomic assignment.

|  | Freshwater sites | | | Marine sites | | | | | |
| --- | --- | --- | --- | --- | --- | --- | --- | --- | --- |
|  | F1 | F2 | F3 | M1 | M2 | M3 | M4 | M5 | M6 |
| Total 16S sequences | 24,611 | 27,187 | 30,622 | 48,675 | 36,194 | 29,723 | 38,295 | 35,753 | 40,100 |
| Classified 16S sequences | 21,301 | 24,786 | 23,292 | 47,035 | 35,494 | 29,054 | 37,175 | 33,576 | 38,267 |
| 16S-ASVs | 1,358 | 1,074 | 1,057 | 467 | 430 | 407 | 396 | 942 | 430 |
| Classified 16S-ASVs | 827 | 692 | 781 | 410 | 389 | 350 | 352 | 676 | 378 |
| Total ITS2 sequences | 100,807 | 108,813 | 126,547 | 83,377 | 89,190 | 75,437 | 83,732 | 74,465 | 90,210 |
| Classified ITS2 sequences | 65,788 | 54,757 | 39,994 | 1,974 | 3,229 | 1,521 | 4,133 | 6,312 | 1,319 |
| ITS2-ASVs | 1,255 | 1,768 | 467 | 197 | 263 | 184 | 252 | 570 | 191 |
| Classified ITS2-ASVs | 1,113 | 1,374 | 209 | 40 | 53 | 38 | 36 | 178 | 25 |

Table S3. List of species associated with environmental variables in our study.

| **Group** | **Species** |
| --- | --- |
| Bacteria | Lentisphaeria |
|  | Firmicutes BRH-c20a |
|  | Hydrogenedentia |
|  | Desulfobacteria |
|  | Deinococci |
|  | Patescibacteria ABY1 |
|  | *Marinimicrobia* (SAR406_clade) |
|  | Desulfitobacteria |
|  | Bacteria Sva0485 |
|  | Thermoanaerobaculia |
| Fungi | Leotiomycetes |
|  | Dacrymycetes |
|  | Monoblepharidomycetes |
|  | Laboulbeniomycetes |
|  | *Sarea difformis* |
|  | Arthoniomycetes |
|  | Olpidiomycota GS18 |
| Bacteria + Fungi | Chytridiomycetes |
|  | Sordariomycetes |
|  | Syntrophia |
|  | *Turneriella* |
|  | *Chloroflexi* KD4-96 |
|  | Marinimicrobia (SAR406_ clade) |
|  | *Truepera* |
|  | Chloroflexi TK10 |
|  | Syntrophobacteria |
